# Cerebral artery segmentation based on magnetization-prepared two rapid acquisition gradient echo multi-contrast images in 7 Tesla magnetic resonance imaging

**DOI:** 10.1101/2019.12.12.870840

**Authors:** Uk-Su Choi, Hirokazu Kawaguchi, Ikuhiro Kida

## Abstract

Cerebral artery segmentation plays an important role in the direct visualization of the human brain to obtain vascular system information. At ultra-high field magnetic resonance imaging, hyperintensity of the cerebral arteries in T1 weighted (T1w) images could be segmented from brain tissues such as gray and white matter. In this study, we propose an automated method to segment the cerebral arteries using multi-contrast images including T1w images of a magnetization-prepared two rapid acquisition gradient echoes (MP2RAGE) sequence at 7 T. The proposed method employed a seed-based region-growing strategy with the following procedures. (1) Two seed regions were defined by Frangi filtering applied to T1w images and by a simple calculation from multi-contrast images, (2) the two seed regions were combined, (3) the combined seed regions were expanded using a region growing algorithm to acquire the cerebral arteries. Time-of-flight (TOF) images were obtained as a reference to evaluate the proposed method. We successfully performed vessel segmentations from T1w MP2RAGE images, which mostly overlapped with the segmentations from the TOF images. As large arteries can affect the normalization of anatomical images to the standard coordinate space in functional and structural studies, we also investigated the effect of the cerebral arteries on spatial transformation using vessel segmentation by the proposed method. As a result, the T1w image removing the cerebral arteries showed better agreement with the standard atlas compared with the T1w image containing the arteries. Thus, because the proposed method using MP2RAGE images can obtain brain tissue anatomical information as well as cerebral artery information without need for additional acquisitions such as of the TOF sequence, it is useful and time saving for medical diagnosis and functional and structural studies.

## 1. Introduction

The visualization of the cerebral vasculature plays an important role in the diagnosis of vascular diseases such as hemorrhage or aneurysms (Stapf et al., 2006; Matsushige et al., 2016) but also to differentiate between anatomical areas in different brain regions (Neumann et al., 2016). Uncertain segmentation of brain tissues and blood vessels can lead to misidentifying the localization of the electroencephalographic signal source and lead to anatomical confounding in certain cortical regions with large vessels in studies of brain function and of cortical characteristics, such as cortical thickness(Fiederer et al., 2016; Viviani, 2016). Therefore, accurate segmentation of blood vessels is essential for neuroimaging research and medical image analysis.

Time-of-flight (TOF) magnetic resonance angiography (MRA) has been widely used as a non-invasive technique to contrast between blood and brain tissues (Lévy et al., 1994). The unsaturated blood flow of cerebral arteries on the TOF image is brighter than the saturated background of other brain tissues. As at ultra-high fields (UHFs), the longitudinal relaxation times (T1) are extended both for blood and brain tissues, leading to enhanced contrast between them, utilization of UHFs for vessel imaging is more efficient (Park et al., 2018). However, there is less anatomical information such as gray matter (GM) and white matter (WM) in TOF images due to background suppression.

Magnetization-prepared rapid gradient echo (MPRAGE) is a candidate for the visualization of blood vessels indicating high signal intensity of arterial blood vessels while the background shows intermediate signal intensities, resulting in high vessel-to-background contrast and is potentially suitable for MRA display (Penumetcha et al., 2008, Wrede et al 2014). Although the T1 weighted (T1w) image in the MPRAGE sequence provides good anatomical information and differentiation of GM and WM, it is difficult to automatically segment the high signal arteries from the intermediate signal tissues in this image. The enhanced contrast between cerebral arteries and brain tissues in the T1w image at UHF can help automatically segment cerebral arteries from the brain (Maderwald et al., 2008; Van de Moortele et al., 2009; Fiederer et al., 2016; Gulban et al., 2018).

Magnetization prepared two rapid acquisition gradient echoes (MP2RAGE) can obtain three different images, a T1 map (T1) and a uniform T1w image with background noise (UNI) and without background noise (UNIDEN) derived by two images at different inversion times, i.e., the first and second inversion gradient echo images (INV1 and INV2, respectively) (Marques et al., 2010). The MP2RAGE sequence has an advantage over the MPRAGE sequence for signal homogeneity in the T1w image (UNI) because the inhomogeneity is cancelled out when the T1w image in the MP2RAGE sequence is calculated from two inversion gradient echo images. As multi-contrast images in the MP2RAGE sequence are useful to segment brain tissues such as GM, WM, and cerebrospinal fluid (CSF) (Choi et al., 2019), the MP2RAGE sequence may also help segment the cerebral arteries.

In this study, we propose an automated method to segment the cerebral arteries using multi-contrast images in the MP2RAGE sequence at 7 T. The proposed method is based on our brain tissue segmentation (MP2rage based RApid SEgmentation; MP2RASE) (Choi et al., 2019) combined with a seed-based approach and a region-growing algorithm. The proposed method using the MP2RAGE sequence was evaluated using vessel segmentation with the TOF sequence as a reference. In addition, we investigated the effect of cerebral arteries on the spatial transformation of T1w images to a standard coordinate space, i.e., the Montreal Neurological Institute (MNI) space, which is ordinarily used in the process of functional and structural MR imaging (MRI) studies.

## 2. Materials and Methods

### 2.1 Subjects

Four volunteers (four men, aged 34–49 years) without history of neurological disease or any other medical condition participated in this study after providing written informed consent. All experiments were approved by the Ethics and Safety Committees of National Institute of Information and Communications Technology.

### 2.2 MRI acquisition

The experiments were performed on a 7-T investigational MRI scanner (MAGNETOM 7T; Siemens Healthineers, Erlangen, Germany) with a 32-channel head coil (Nova Medical, Wilmington, MA). The MP2RAGE was acquired using a research sequence from Siemens Healthineers and using the following parameters: repetition time = 5000 ms, echo time = 3.43 ms, inversion times = 800 ms/2600 ms, flip angles = 4°/5°, matrix = 368 × 368 × 256, voxel size = 0.7 × 0.7 × 0.7 mm^3^, GRAPPA acceleration factor = 3, and scan time = 9 min 27 s (Choi et al., 2019). To evaluate the proposed method, we additionally acquired 3D TOF images with the following parameters: voxel resolution = 0.3 × 0.3 × 0.4 mm^3^, repetition time = 20 ms, echo time = 4.47 ms, flip angle = 18°, matrix = 704 × 704 × 192, GRAPPA acceleration factor = 4, and scan time = 12 min 19 s (Wrede et al., 2014; Matsushige et al., 2016). The acquisition region of the TOF sequence included large cerebral arteries such as the anterior cerebral artery (ACA), middle cerebral artery (MCA), and internal carotid artery (ICA).

### 2.3 Vessel segmentation using MP2RAGE images

The image processing pipeline in the proposed method is shown in Fig. 1. All MP2RAGE images were stripped to remove non-brain structures using the BET toolbox in FSL 5.0.10 (FMRIB, Oxford, UK) during the brain segmentation procedure. After removing non-brain structures, we normalized the intensities of all images for the brain tissue segmentation using a feature-scaling method described in Eq. 1 because MP2RAGE images have different intensity scales. S_raw_ indicated raw intensities, min (S_raw_) indicated the minimum intensity, and max (S_raw_) indicated the maximum intensity of each MP2RAGE image. 

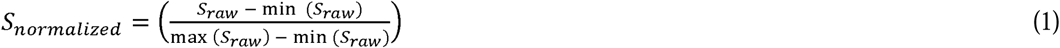

**Figure 1.**
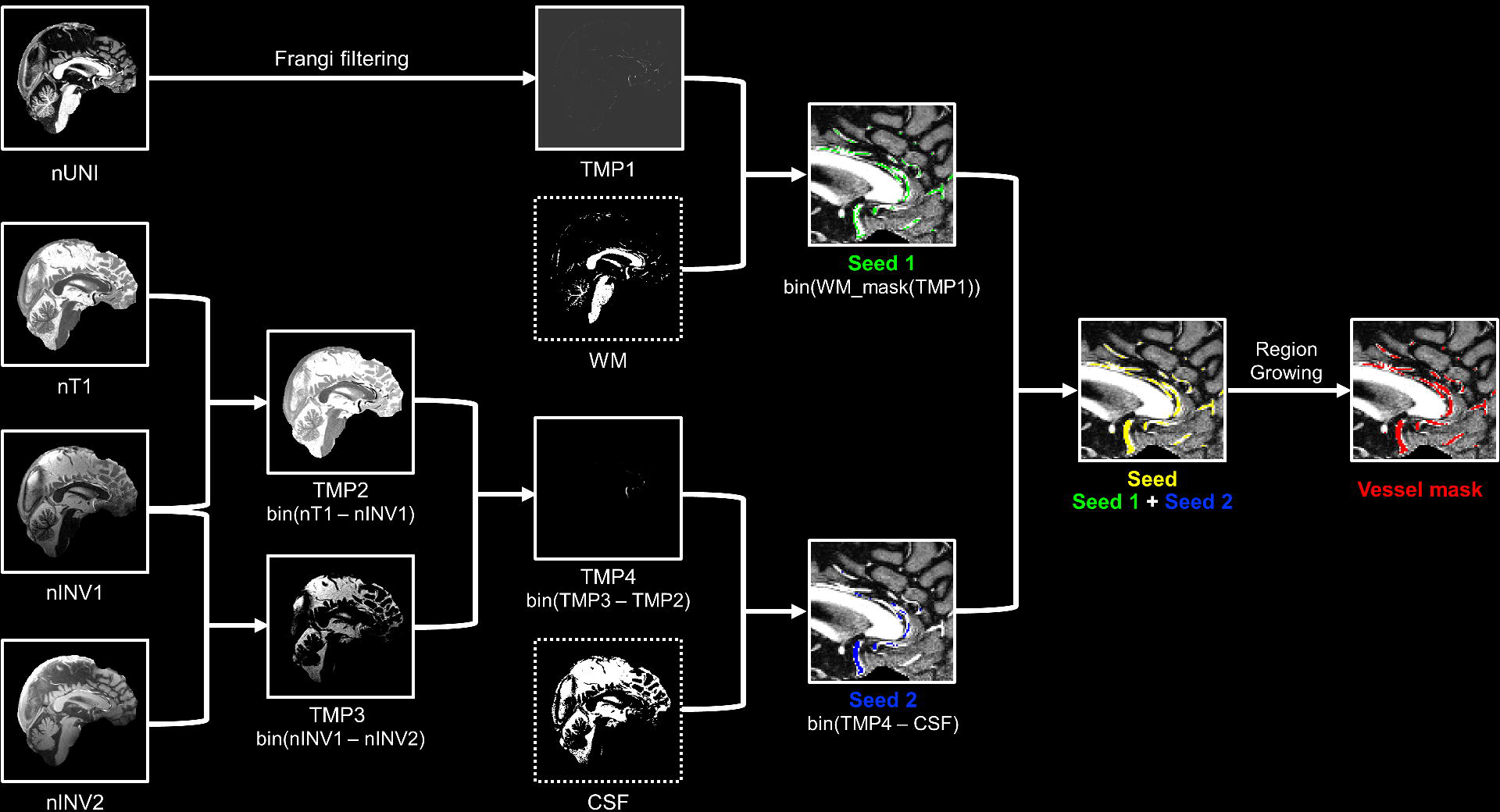
Image processing pipeline of the proposed method. Dotted squared images were constructed from the brain tissue segmentation method (Choi et al., 2019). CSF: cerebral spinal fluid, nINV1: normalized 1^st^ inversion time image, nINV2: normalized 2^nd^ inversion time image, nT1: normalized T1 map, nUNI: normalized uniform T1w image, WM: white matter.

First, we employed a Frangi filter (Frangi et al., 1998), which has been used for vessel segmentation in TOF (Hsu et al., 2019) and MPRAGE (Fiederer et al., 2016; Oliveira et al., 2016;), to construct a binary vessel mask (Seed 1 in Fig. 1) from the normalized UNI (nUNI) image (Kroon, 2009). The filtering was performed with the following parameters; α = 0.5, β = 0.5, γ = 5.0 (Antiga, 2007). The filtered image was binarized with a threshold of a 10% robust range of non-zero voxels and masked out by a WM binary mask, which was segmented using our previous method (Choi et al., 2019). Second, a binary vessel mask (Seed 2 in Fig. 1) was constructed using three normalized contrast images (nINV1, nINV2, and nT1) and a CSF binary mask, which was also segmented using our previous method (Choi et al., 2019). The binary vessel masks, i.e., seed 1 and seed 2, were combined for the region-growing procedure. Finally, the vessel mask was obtained by expanding the voxels of the combined binary vessels masks by a region growing procedure using our in-house MATLAB code including a recursive region growing algorithm (Daniel, 2011).

### 2.4 Vessel segmentation using TOF images

The intensities of the TOF images were normalized using the feature-scaling method described in Eq. 1. The vessels were segmented from the normalized TOF images using a Frangi filter with the following parameters; α = 0.5, β = 0.5, γ = 5.0. The filtered images were binarized with a threshold of a 25% robust range of non-zero voxels to construct the TOF binary vessel mask.

### 2.5 Comparison between vessel segmentations using MP2RAGE and TOF images

We employed the Dice coefficient to evaluate the similarity of the vessel segmentations of MP2RAGE images with those of TOF images as a reference. Before evaluating the similarity, the two datasets were coregistered because the dimensions between TOF and MP2RAGE images differ (Supplementary Fig. 1). TOF images were initially registered to the UNIDEN of the MP2RAGE sequence to acquire the transformation matrix using FLIRT in FSL. The matrix was applied to binary vessel masks from TOF images with nearest-neighbor interpolation. Next, the pre-registered vessel masks from the TOF images were registered to vessel masks from MP2RAGE for fine spatial transformation using the non-linear registration method of Advanced Normalization Tools (ANTs) (Avants et al., 2011) with nearest-neighbor interpolation. After registration, we defined two fields of view (FOVs) for the evaluation with the Dice coefficient. One FOV was the acquisition region of the TOF sequence, which was fully covered by the MP2RAGE image (Supplementary Fig. 1). Another FOV was manually defined including large arteries such as the ACA, MCA, and ICA as local FOV regions. The vessel segmentations within two FOVs were evaluated to calculate the Dice coefficient with Eq. 2. 

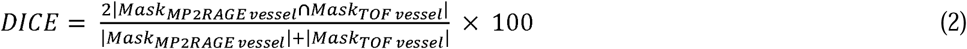

### 2.5. Spatial normalization to the standard space

Cerebral arteries in the UNIDEN T1w image were firstly removed based on vessel segmentation by the proposed method using an in-house script. The T1w images with and without cerebral arteries were spatially normalized to the MNI152 template in FSL using an in-house script including ANTs.

## Results

### 3.1 Vessel segmentation using MP2RAGE images

Two binary vessel masks were constructed by independent processes: Frangi filtering of the nUNI image and a simple calculation of MP2RAGE images. The masks partially overlapped, indicating that the two seed regions represented independent parts of the cerebral arteries (Fig. 2). Most voxels in the arteries were extracted as a vessel mask by Frangi filtering (Fig. 2a) but some voxels in relatively larger arteries were not obtained. Conversely, the voxels, which were not detected by Frangi filtering, were represented as a vessel mask using a simple calculation of MP2RAGE images (Fig. 2b). Application of a region-growing algorithm for a combination of seed 1 and seed 2 extracted most voxels in the arteries as a vessel mask (Fig. 2c), which shown in 3D reconstruction view of the cerebral arteries (Fig. 3). Large arteries such as the MCA, ACA, and ICA were well segmented.

**Figure 2.**
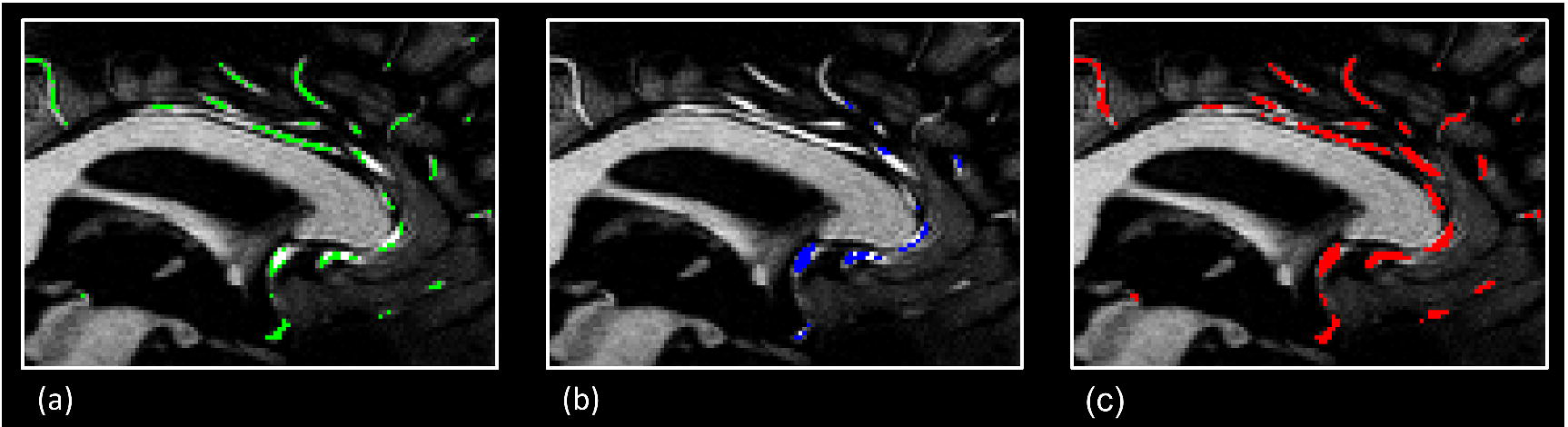
Visual presentation of segmentation at major stages. (a) segmentation by Frangi filtering applied to the T1w image, (b) segmentation by a simple calculation of multi-contrast images, (c) segmentation in the final result of the proposed method. Segmentation masks were overlaid to the T1w image, which indicated a sagittal view of regions around the anterior cerebral artery.

**Figure 3.**
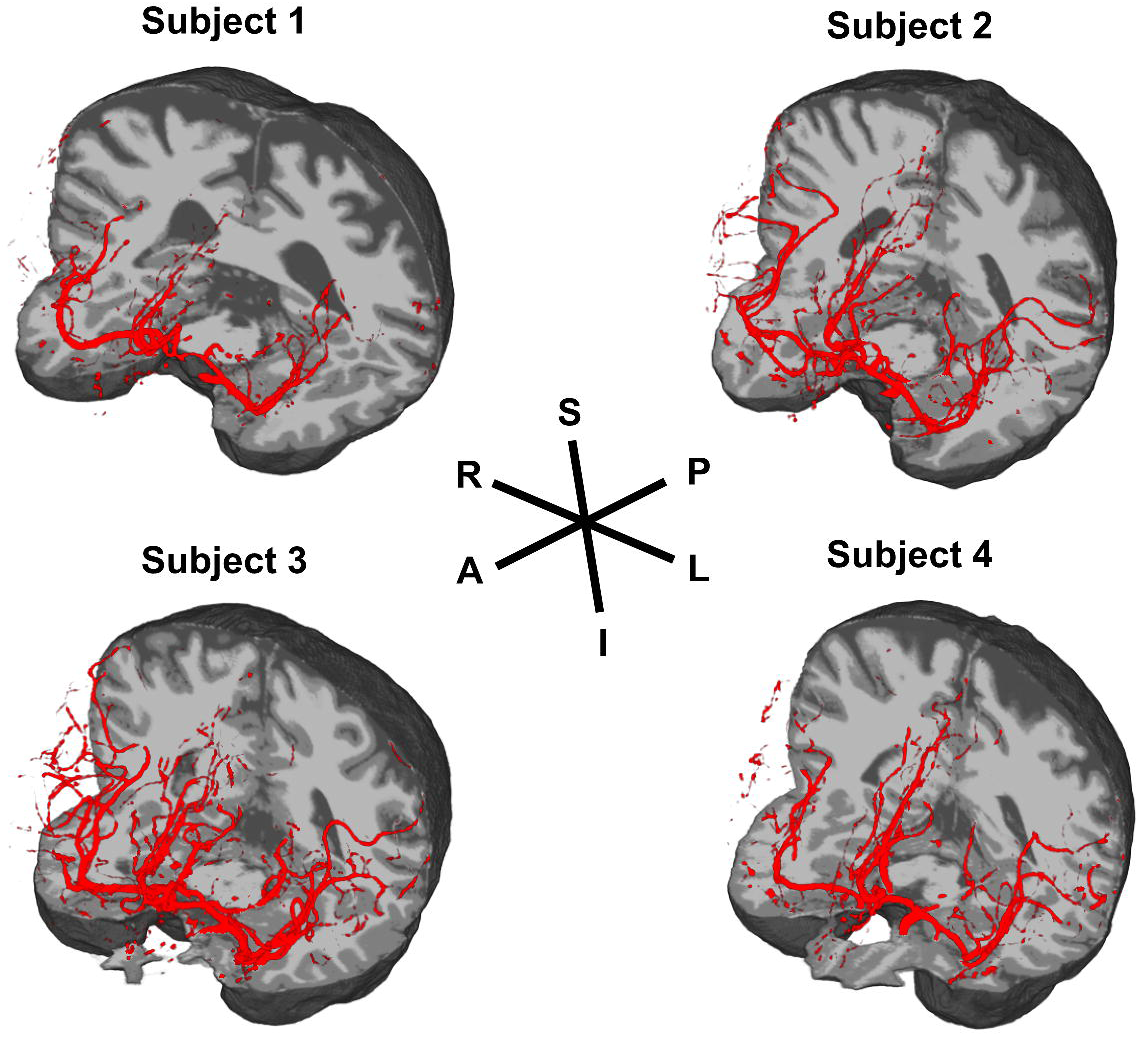
3D visualization of vessel segmentation by the proposed method in all subjects.

### 3.2 Comparison between vessel segmentations of MP2RAGE and TOF images

Figure 4 shows a comparison of vessel segmentations from MP2RAGE images by the proposed method and from TOF images by Frangi filtering in 3D reconstruction view. Most arteries extracted from the MP2RAGE images by the proposed method showed good agreement with the ones extracted from the TOF images. Larger arteries such as the MCA, ACA, and ICA well overlapped, but peripheral branches of arteries were mismatched between vessel segmentations of MP2RAGE and TOF images. We evaluated the proposed method by comparison to TOF vessel segmentation using the Dice coefficient for whole FOV regions, which overlapped between the two sequences, and the local FOV regions including larger arteries (Fig. 5). The Dice coefficients with vessel segmentation from TOF images were approximately 40% for the whole FOV regions and 55% for local regions when vessels in the T1w MP2RAGE image were segmented by Frangi filtering (seed 1), which was the same method as that used in TOF segmentation. The coefficients by simple calculation from the MP2RAGE images (seed 2) for both regions were almost the same as those obtained with Frangi filtering. After combining the two seeds, the similarities of vessel segmentation dramatically improved from 40 to 57% for the whole FOV regions and from 55 to 73% for the local FOV regions. The Dice coefficients showed a lower similarity for the whole FOV region because this approach may not be able to properly handle the small branches of large cerebral arteries.

**Figure 4.**
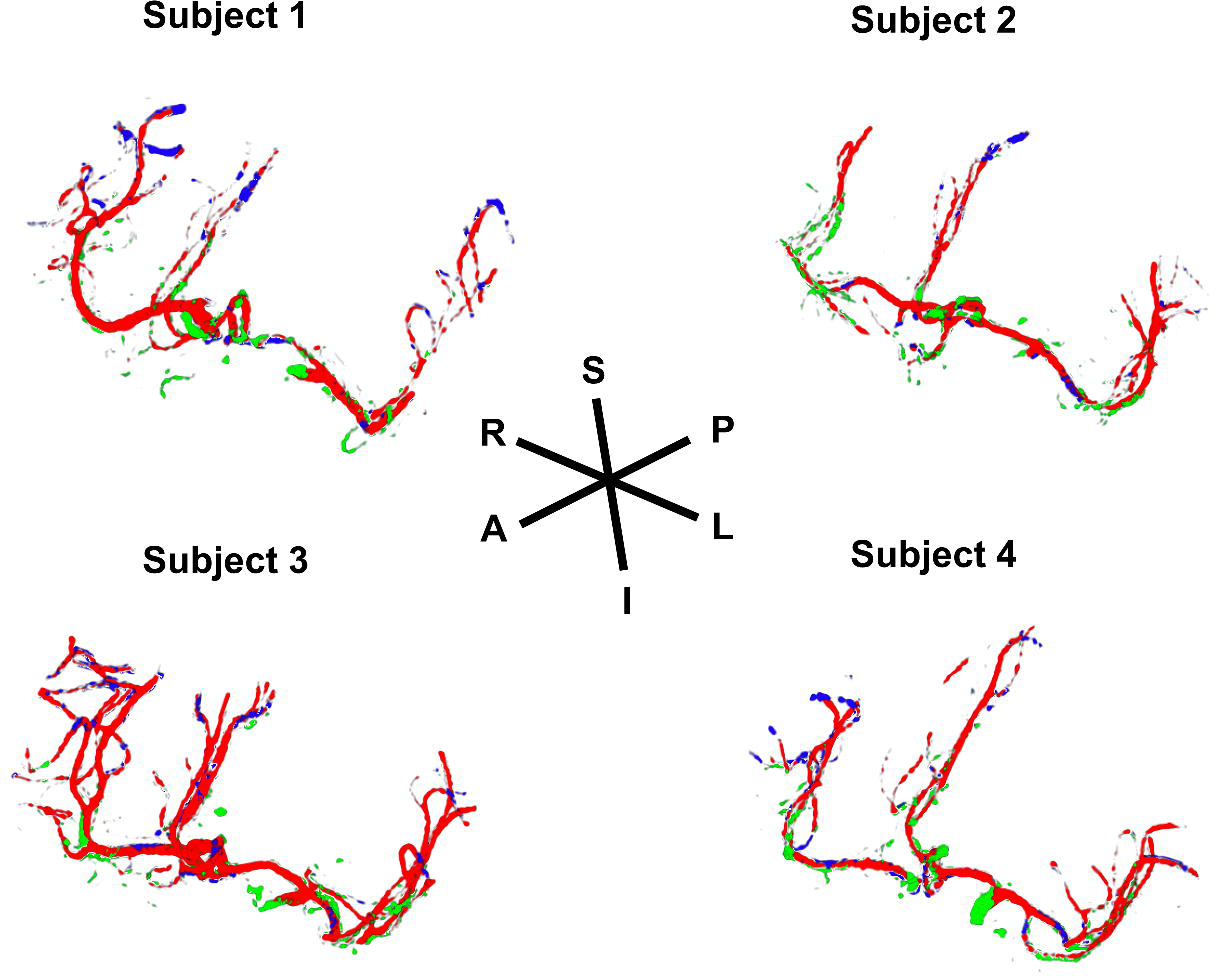
Comparison of cerebral artery segmentation between the results from MP2RAGE and TOF images in all subjects. Red color: overlapped segmentation between two methods, Green color: segmentation only from MP2RAGE images, Blue color: segmentation only from TOF images.

**Figure 5.**
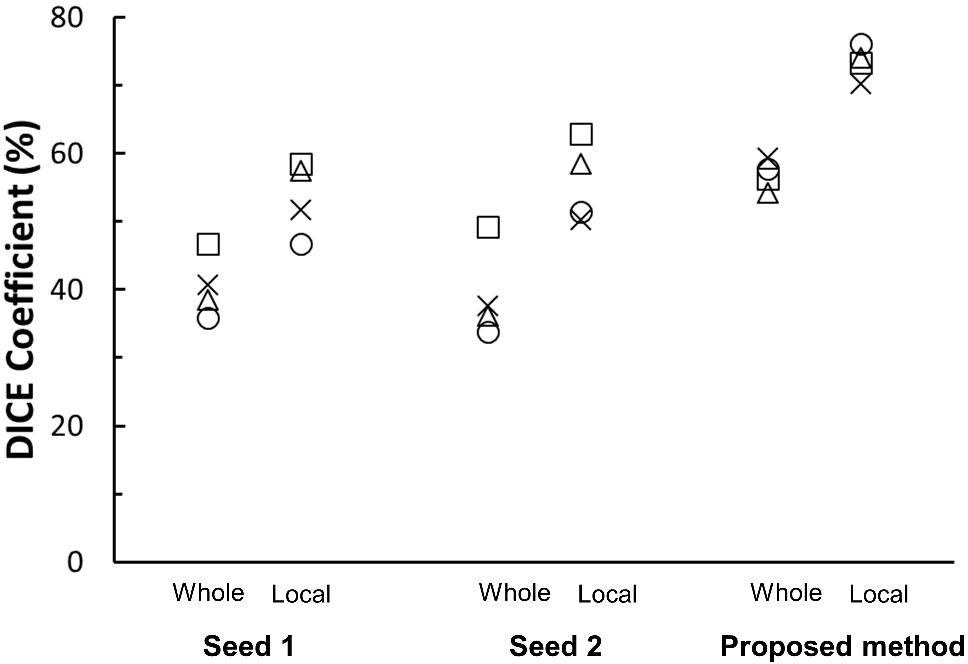
Similarity between vessel segmentations at major stages from MP2RAGE and TOF images. Seed 1: segmentation by Frangi filtering, Seed 2: segmentation by a simple calculation of MP2RAGE images. The Dice values show vessel segmentation from the whole FOV region (whole) and from large vessel FOV regions (local) in all subjects.

### 3.3 Influence of cerebral arteries on normalization to standard coordinate space

We examined the influence of the cerebral arteries on the spatial transformation of T1w images to a standard coordinate space, i.e., the MNI space, which is ordinarily used in the process of functional MRI and voxel-based morphometry (VBM). Figure 6 shows the spatial transformation in the T1w image with and without cerebral arteries, segmented by the proposed method. By removing the MCA and ACA from the T1w image, the GM intensities in the frontal orbital cortex and cingulate cortex recovered when the T1w image was normalized to the standard coordinate space. Thus, the T1w image after normalization excluding the cerebral arteries is very consistent with the MNI template in FSL.

**Figure 6.**
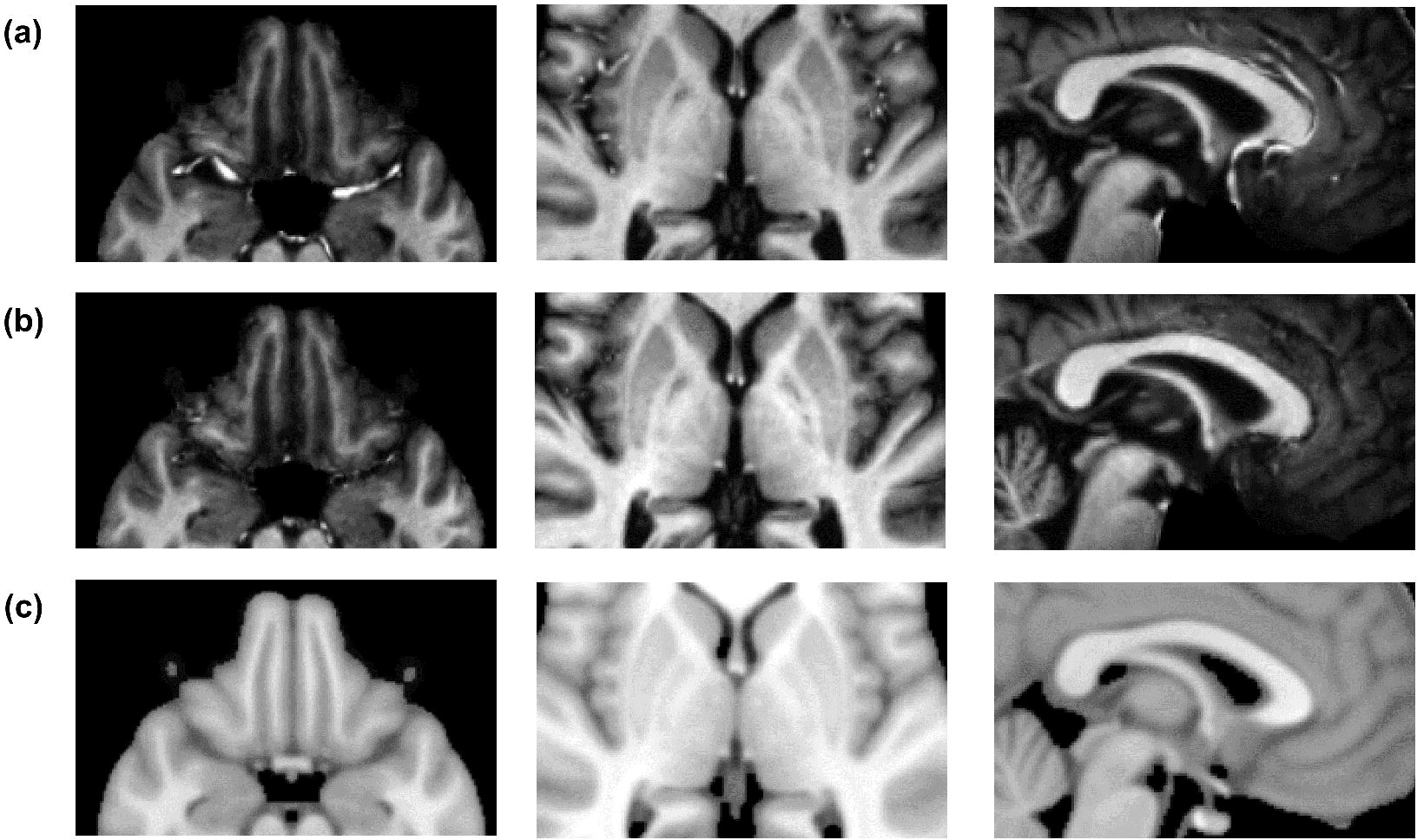
Effect of the cerebral arteries on normalization to the MNI coordinates. a) Normalized T1w image with cerebral arteries, (b) Normalized T1w image without cerebral arteries, (c) MNI template. The 1^st^, 2^nd^, and 3^rd^ columns from the left side indicate the axial view of the orbitofrontal cortex and insular cortex and the sagittal view of the cingulate cortex, respectively. The MCA in the 1^st^ and 2^nd^ columns and the ACA in the 3^rd^ column are shown in (a) but not in (b).

### Discussion

We developed a novel method to automatically segment cerebral arteries with a region growing algorithm for two combined seed masks from T1w MP2RAGE sequence images at 7 T. The two seed masks were defined by Frangi filtering of the T1w image and by a simple calculation of multi-contrast images in the MP2RAGE sequence. The seed mask of the region growing algorithm, which has been widely used as a segmentation method for medical images (Malek et al., 2012), is important to define core voxels of arteries for the robust region growth of cerebral arteries. Most of the seed voxels were defined by Frangi filtering, which has been employed to segment blood vessels in the retina (Oliveira et al., 2016) and the brain (Fiederer et al., 2016; Hsu et al., 2019). The filtering can be useful to differentiate blood vessels from brain tissues in high contrast images, i.e., TOF images but not with less suppressed tissue images such as T1w image (i.e., UNI) in the MP2RAGE sequence. The advantage of multiple contrasts for segmentation in MP2RAGE images has been described in previous studies (Wang et al., 2018; Choi et al., 2019). Therefore, as more seed voxels might be needed for segmentation, we additionally defined the seed voxels using a simple calculation of multi-contrast images from the MP2RAGE sequence, which provided better results.

As the major cerebral arteries are located in frontal parts of the brain such as the anterior cingulate, orbitofrontal, and insular cortices, which are core brain regions in cognitive studies (Bechara et al., 2000; Bush et al., 2000; Mutschler et al., 2009), the large cerebral arteries might influence the standard space transformation of T1w images. Better agreement with the MNI atlas was achieved when the T1w image was transformed to exclude the cerebral arteries using vessels mask extracted by the proposed method compared with the agreement of T1w images that contained the cerebral arteries (Fig. 6), which led to more precise transformation evaluation. Precise transformation information could be important to assess the thickness of the cerebral cortex and volume using methods such as VBM (Kotikalapudi et al., 2019) and to obtain accurate location of neuronal activation such as olfactory functional MRI study (Gottfried et al., 2002). Especially, as recent UHF studies have focused on fine brain structures and brain function at the submillimeter scale without spatial smoothing, functional images with high spatial resolution can be affected because cerebral arteries have the potential to induce errors of structural transformation in those cortical areas, which can induce mis-localization of specific brain function.

The proposed method has limitations to be addressed in further studies. The proposed method could reliably detect major cerebral arteries such as the MCA, ACA, and ICA but failed to segment small peripheral vessels possibly due to less contrast with the surrounding brain tissues because of limited resolution, slow blood flow rate, and impaired image quality involving distortion or inhomogeneity. B1 mapping of the SA2RAGE sequence (Eggenschwiler et al., 2012) could possibly improve the inhomogeneity of the T1w image at UHFs. The enhanced contrast to noise ratio between GM and WM could also help improve the segmentation by changing the acquisition MR parameters of the MP2RAGE sequence such as the inversion times and flip angles (Tanner et al., 2012; Marques and Gruetter, 2013). Although increased spatial resolution could be another possibility to measure peripheral arteries, a low signal to noise ratio and longer scan time comprise trade-offs for the MP2RAGE sequence.

We employed vessel segmentation of TOF images with Frangi filtering as reference to compare with the proposed method based on the Dice coefficient. The quality of vessel segmentation in both TOF and the proposed method depends on the parameter and threshold of Frangi filtering, which also depends on the quality of the TOF (Phellan et al., 2017). In fact, parts of major arteries were not segmented using TOF images with Frangi filtering. Further evaluation of the proposed method will be needed by a quantitative verification with the different acquisition and analysis parameters (Frangi filtering and threshold), which may lead to further improvement.

In conclusion, we proposed a novel method to automatically segment cerebral arteries using MP2RAGE images at 7 T. As we previously developed a method to segment brain tissues using the MP2RAGE sequence (MP2RASE) (Choi et al., 2019), together, a single MP2RAGE sequence acquisition could provide automated brain tissue masks of GM, WM, and CSF as well as of blood vessels. In addition, the proposed method could be used to derive accurate spatial transformation information with reference to the standard coordinate space by removing the influences of cerebral arteries from T1w images. Thus, because the proposed method of vessel segmentation using MP2RAGE sequence can obtain brain tissue anatomical information as well as cerebral artery information without any additional acquisitions such as TOF sequence, it is useful and time saving for medical diagnosis and functional and structural studies.

## Acknowledgements

We would like to thank Dr. Kober for the development of MP2RAGE sequence and technical supports. This work was supported by the JSPS KAKENHI [grant number JP18K07701, JP18H04084, and JP19H03537] from the Japanese Ministry of Education, Culture, Sports, Science and Technology (IK).

## Abbreviations

ACA: anterior cerebral artery
CSF: cerebrospinal fluid
FOV: field of view
GM: gray matter
ICA: internal carotid artery
INV1: first inversion gradient echo image
INV2: second inversion gradient echo image
MCA: middle cerebral artery
MNI: Montreal Neurological Institute
MRA: magnetic resonance angiography
MPRAGE: Magnetization-prepared rapid gradient-echo
MP2RAGE: magnetization-prepared two rapid acquisition gradient echoes
TOF: time-of-flight
UHF: ultra-high field
UNI: uniform T1w image with background noise
UNIDEN: uniform T1w image without background noises
VBM: voxel-based morphometry
WM: white matter

## Figure caption

**Supplementary Figure 1.**
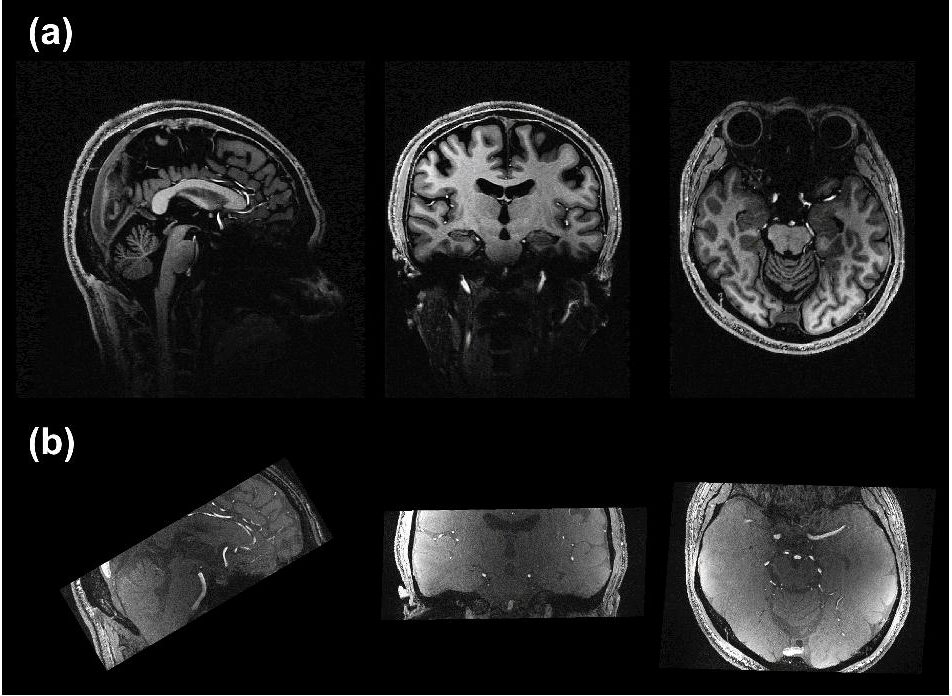
MP2RAGE and TOF images. (a) uniform T1w image without background noise from the MP2RAGE sequence, (b) TOF image.

